# Functional Territories of Human Dentate Nucleus

**DOI:** 10.1101/608620

**Authors:** Xavier Guell, Anila M D’Mello, Nicholas A Hubbard, Rachel R Romeo, John DE Gabrieli, Susan Whitfield-Gabrieli, Jeremy D Schmahmann, Sheeba Arnold Anteraper

## Abstract

Anatomical connections link the cerebellar cortex with multiple distinct sensory, motor, association, and paralimbic areas of the cerebrum. These projections allow a topographically precise cerebellar modulation of multiple domains of neurological function, and underscore the relevance of the cerebellum for the pathophysiology of numerous disorders in neurology and psychiatry. The majority of fibers that exit the cerebellar cortex synapse in the dentate nuclei (DN) before reaching extracerebellar structures such as cerebral cortex. Although the DN have a central position in the anatomy of the cerebello-cerebral circuits, the functional neuroanatomy of human DN remains largely unmapped. Neuroimaging research has redefined broad categories of functional division in the human brain showing that primary processing, attentional (task positive) processing, and default-mode (task negative) processing are three central poles of neural macro-scale functional organization. This new macro-scale understanding of the range and poles of brain function has revealed that a broad spectrum of human neural processing categories (primary, task positive, task negative) is represented not only in the cerebral cortex, but also in the thalamus, striatum, and cerebellar cortex. Whether functional organization in DN obeys a similar set of macroscale divisions, and whether DN are yet another compartment of representation of a broad spectrum of human neural processing categories, remains unknown. Here we show for the first time that human DN is optimally divided into three functional territories as indexed by high spatio-temporal resolution resting-state MRI in 60 healthy adolescents, and that these three distinct territories contribute uniquely to default-mode, salience-motor, and visual brain networks. These conclusions are supported by novel analytical strategies in human studies of DN organization, including 64-channel MRI imaging, data-driven methods, and replication in an independent sample. Our findings provide a systems neuroscience substrate for cerebellar output to influence multiple broad categories of neural control - namely default-mode, attentional, and multiple unimodal streams of information processing including motor and visual. They also provide a validated data-driven mapping of functions in human DN, crucial for the design of methodology and interpretation of results in future neuroimaging studies of brain function and dysfunction.

## INTRODUCTION

Neuroimaging research has redefined broad categories of functional division in the human brain. Specifically, it has shown that (i) primary processing, (ii) task-positive non-primary processing, and (iii) task-negative (default-mode) processing are three central poles of neural macro-scale functional organization (Fox et al. 2005, Greicius et al. 2003, Margulies et al. 2016). Primary processing includes motor, somatosensory, auditory, and visual systems. Non-primary task-positive processing includes top-down goal-directed systems active in attention demanding activities such as working memory tasks, and bottom-up stimulus-driven systems that direct attention towards salient stimuli (Corbetta & Shulman 2002, Fox et al. 2006, Seeley et al. 2007). Task-negative regions are engaged in abstract higher-order association processes that are furthest removed from primary systems (Margulies et al. 2016, Mesulam 1998) and anti-correlated with task-positive networks (Fox et al. 2005, Zhou et al. 2018), including unfocused processes with low attentional demands such as mind wandering, and related states of autobiographical memory retrieval and introspection (Buckner et al. 2008, Greicius et al. 2003).

This new macro-scale understanding of the range and poles of brain function has revealed that not only the cerebral cortex (Yeo et al. 2011), but also the thalamus (Hwang et al. 2017), striatum (Choi et al. 2012), and cerebellar cortex (Buckner et al. 2011) contribute to a broad spectrum of human neural processing categories. Specifically, primary processing networks such as somatomotor, task-positive networks such as ventral and dorsal attention, and task-negative default mode network are all represented within each of these structures.

Whether the dentate nuclei (DN) of the cerebellum obey a similar set of macroscale functional divisions, and whether the DN are yet another compartment of broad-spectrum representation of human brain function, remains unknown. DN viral tracing studies have described a motor versus non-motor dichotomy (Dum & Strick 2003), a bi-modal division that has been echoed in human resting-state fMRI (Bernard et al. 2014), task-based fMRI (Küper et al. 2011, 2012, 2014; Thürling et al. 2011), and diffusion tensor imaging (DTI; Steele et al. 2017). Anatomical and neuroimaging investigations indicate that the central components of functional specialization in human DN may extend beyond a dual motor versus non-motor classification, and span primary, task-positive, and task-negative domains of brain function. First, these systems are all represented in cerebellar cortex (Buckner et al. 2011, Guell et al. 2018b), in close correspondence with cerebral cortical organization (Margulies et al. 2016, Yeo et al. 2011). Second, the DN are a central node in the anatomy of cerebellar connections, relaying the majority of fibers exiting the cerebellar cortex and participating in multiple reverberating systems within the cerebellar circuits. The DN are the largest and most lateral of the cerebellar nuclei, and receive the majority of cerebellar cortical efferents (Haines & Dietrichs 2011). Ascending projections of the dentate nuclei are directed principally to the thalamus, connecting cerebellar cortex to thalamo-cortical projections and thus to sensorimotor cortices and higher-order association areas. Multiple reverberating patterns exist in the connectivity between cerebellar cortex, DN, and extracerebellar structures. First, neurons in DN project back to cerebellar cortical areas from which they receive input (Dietrichs 1981). Second, each inferior olivary nucleus (i) projects directly to a specific territory of cerebellar cortex, (ii) projects to the DN territory that receives projections from that particular territory of cerebellar cortex, (iii) and at the same time receives projections from that particular DN territory (as reviewed in De Zeeuw et al. 1998). Third, current anatomical evidence points to reciprocal reverberating loops between defined areas of cerebral cortex, pons, cerebellar cortex, dentate, and thalamus. That is, a cerebral cortical area that sends projections to the cerebellar cortex receives feedback from that same cerebellar corticonuclear microcomplex (Kelly & Strick 2003). This rich set of connections provides an anatomical basis for the hypothesis that a broad range of macro-scale functional categories in the brain are represented in DN.

Gradient-based and data-driven clustering analyses of resting-state data, a methodology that remains largely unexplored in DN, can provide critical information necessary to test this hypothesis. The identification of the main functional subdivisions of human DN requires a data-driven approach independent from a-priori assumptions regarding the nature and localization of functional domains, and flexible to include as many components as necessary to fully identify the principal poles and spectrum of DN specialization.

Here we set out to identify for the first time the central components of functional organization in human DN. We used gradient-based and data-driven clustering methods that have previously unmasked these central aspects of functional neuroanatomy in cerebral (Margulies et al. 2016) and cerebellar cortex (Guell et al. 2018b). 64-channel resting-state MRI data combined with simultaneous multi-slice acquisition provided unprecedented DN spatial and temporal resolution useful to overcome technical difficulties of functional mapping in small brain territories. We then attempted to replicate our results in a separate set of participants, validating our findings and their utility for future independent human studies of DN.

## METHODS

### Study Participants

60 right-handed participants were included in this study. 20 participants (10 male, mean age = 13.64, age range = 12-14) were included in the discovery sample, and 40 participants (18 male, mean age = 15.13, age rage = 14-16) were included in the replication sample. A comment regarding participants’ age ranges is included in the last two sections of the discussion. These participants were recruited as controls subjects as part of two ongoing larger studies (one study examined brain correlates of depression and anxiety in adolescents, and another study investigated brain correlates of cognitive processing, social processing, socioeconomic status, and academic achievement in adolescents). Written informed consent was collected from all participants in accordance with guidelines established by the Massachusetts Institute of Technology Committee on the Use of Humans as Experimental Subjects (discovery sample) and the Partners Health Care Institutional Review Board (replication sample). None of the participants had a self-reported or family-reported history of psychiatric or neurological illness.

### MRI structural and resting-state data

Imaging data for the discovery and replication samples were collected on two distinct 3T Siemens PRISMA MRI scanners with vendor-provided 64 Channel head coil (Siemens Healthcare, Erlangen, Germany). Both scanners corresponded to the same model and all participants within each sample were scanned in the same scanner. Scanning parameters were identical for both the discovery and replication samples. High-resolution structural data (0.8 mm isotropic voxels) were acquired using a T1-weighted MPRAGE sequence with duration 7 minutes 50s (in-plane acceleration factor of 2). Scan parameters for TR/TE/TI/Flip Angle were 2.4s/2.18ms/1.04s/8°. Anatomical scans were immediately followed by resting-state scans, during which subjects were asked to stay awake and keep their eyes fixated on a cross hair. Two resting-state sessions per participant were acquired for the discovery sample, and four sessions per participant were acquired for the replication sample. Scan parameters (T2*-weighted EPI sequence) for TR/TE/Flip Angle/echo spacing/bandwidth were 800ms/37ms/85°/0.58ms/2290Hz-per-pixel. 72 interleaved (ascending/foot-head) slices were collected in AC-PC plane using an auto-align procedure to minimize inter-subject variability in data acquisition. Combination of 64Ch array coil and simultaneous multi-slice (SMS) acquisition (multiband factor of 8) (Setsompop et al. 2012) provided high temporal sampling (420 time points during an acquisition window of 5 minutes and 46s for each session in the discovery and replication samples) and spatial resolution (2mm isotropic) while maintaining whole-brain coverage (including the entire cerebellum). Of note, 64Ch array coils tolerate relatively high multi-band factors (8 in the present study) providing improved signal-to-noise ratio (SNR) for all brain areas as demonstrated previously (Keil et al. 2013). Further, unlike in-plane acceleration, SMS imaging strategies to increase temporal sampling have no signal-to-noise penalty for multi-band acceleration (Setsompop et al. 2012).

### Data analysis: preprocessing and calculation of connectivity matrix from dentate nuclei to cerebral cortex

EPI data were realigned, normalized to MNI template, and spatially smoothed with a 4-mm FWHM Gaussian kernel using SPM12 (Wellcome Department of Imaging Neuroscience, London, United Kingdom; www.fil.ion.ucl.ac.uk/spm). Structural images were segmented into white matter, grey matter, and cerebrospinal fluid using SPM12 (Ashburner & Friston 2005). CONN Toolbox (Whitfield-Gabrieli & Nieto-Castanon 2012) was used for calculating connectivity from DN to the whole brain, but only connectivity data from DN to cerebral cortex was used in our analyses. In the discovery sample, for each voxel within DN (as defined using the SUIT DN mask (Diedrichsen et al. 2011)), a 2mm-sphere seed was generated and used as a region of interest for seed-to-voxel whole-brain analysis. Band pass filtering was executed at 0.008-0.09 Hz. The Artifact Detection Toolbox (http://www.nitrc.org/projects/artifact_detect) was used for denoising, as follows. Time points with mean signal intensity outside 3 standard deviations from global mean, and 0.5 mm scan-to-scan motion were flagged as invalid scans and were regressed out along with the six realignment parameters and physiological sources of noise (i.e., three Principal Components of white matter, and three Principal components of cerebrospinal fluid segments, using aCompCor (Behzadi et al. 2007)). Whole brain correlation maps derived from denoised time-series from each voxel in DN for each participant in the discovery sample were used for functional gradient calculations, as follows.

### Data analysis: calculation of functional gradients

Based on novel analytical strategies introduced by Coifman and colleagues (Coifman et al. 2005), functional gradient analyses of resting-state data have unmasked the principal poles and components of functional neuroanatomy in previous investigations of cerebral cortex (Margulies et al. 2016) and cerebellar cortex (Guell et al. 2018b). Here we applied this technique for the first time to the study of resting-state data in DN. In brief, this methodology analyses the similarity between the connectivity patterns of each datapoint within a given brain structure, and extracts functional gradients that represent the principal poles and transitions of connectivity patterns within that structure. More specifically, our analysis started by constructing a connectivity matrix including resting-state correlation values from each voxel in DN to each voxel in cerebral cortex (as described in the previous section). The connectivity pattern of each DN voxel was thus represented as an n-dimensional vector, where n corresponded to the total number of voxels in the cerebral cortex. Because all DN voxels were represented in the same n-dimensional space, cosine distance between each pair of vectors could be calculated, and an affinity matrix was constructed as (1 - cosine distance) for each pair of vectors. This affinity matrix represented the similarity of connectivity patterns for each pair of DN voxels. A Markov chain was then constructed using information from the affinity matrix; information from the affinity matrix was thus used to represent the probability of transition between each pair of vectors. In this way, there was higher transition probability between pairs of DN voxels with similar connectivity patterns. This probability of transition between each pair of DN voxels was analyzed as a symmetric transformation matrix, allowing the calculation of eigenvectors. Eigenvectors derived from this transformation matrix represented the principal orthogonal directions of transition between all pairs of voxels in DN. Diffusion map embedding using functional connectivity values from DN to cerebral cortex thus captured the principal functional gradients of DN functional neuroanatomy. See https://github.com/satra/mapalign for an implementation of this methodology, and our online repository for the application of this methodology to our data (https://github.com/xaviergp/dentate).

### Data analysis: clustering of functional gradients

Functional gradient values are a continuous measure useful for establishing groups (clusters) of DN voxels based on functional similarity. As functional gradients capture relevant dimensions of DN functional neuroanatomy, groups of voxels in DN can be clustered together based on the similarity of their functional gradient values. In this way, the optimal number of functional divisions of DN, and the location of these principal poles and components of DN functional neuroanatomy, can be identified.

We used data-driven methods to establish clustering analysis parameters, including the number of functional gradients to include in the clustering analysis, and the number of clusters that the clustering analysis should detect, as follows. First, eigenvalues of each functional gradient were used to establish the data variability explained by each component, and thus to establish the set of functional gradients (g) that should be included in the clustering analysis. Only those functional gradients explaining a relevant portion of data variability (as determined when visualizing the curve of percentage variability explained by each functional gradient) were considered to represent a relevant aspect of DN functional neuroanatomy, and thus included as dimensions for the clustering analysis. Then, silhouette coefficient analysis was used to identify the optimal number of clusters (c) to be computed for that set of functional gradients (g) (Hastie, Trevor, Tibshirani, Robert, Friedman 2009). Using scikit-learn (Pedregosa & Varoquaux 2011), k-means separated DN voxels in c clusters, minimizing the sum of the squared differences of each data point from the mean within each cluster in a g-dimensional functional space.

Because clustering took place in functional (g) rather than anatomical space, clusters identified were not necessarily composed of contiguous voxels in DN anatomical space. To reduce noise in the spatial distribution of our final DN parcellation, an anatomical cluster-size threshold was used so that only anatomical clusters containing 10 or more contiguous voxels were included in our final definition of functional territories in DN.

### Data analysis: post-hoc characterization of connectivity from DN clusters and validation

To characterize the functional significance of each identified functional territory in DN, each DN functional territory was used as a seed in a seed-to-voxel resting-state functional connectivity analysis. Using the CONN Toolbox (Whitfield-Gabrieli & Nieto-Castanon 2012), Pearson’s correlation coefficients were computed using the denoised time courses, and then converted to normally distributed z-scores using Fisher transformation to allow second-level General Linear Model analyses. To test the possibility that each cluster in DN represented a functional domain that was different from the functional domain of other clusters (unique effect), seed-to-voxel analyses were also calculated by computing the effect of each functional territory against the effect of all other functional territories. The same analyses were performed in the independent replication dataset. Functional connectivity from each functional territory was first visualized after thresholding at the 95^th^ percentile of r median correlation values in the discovery sample – namely, using the same thresholding and sample that was used when calculating functional gradients. We then performed unique effect analyses that were based on statistical significance thresholding, including p<0.001 at the voxel level with a p<0.05 FDR cluster-size correction, in both the discovery and replication samples. An additional lower threshold was also used in the replication sample (p<0.005 at the voxel level with a p<0.05 FDR cluster-size correction) to allow better visualization of brain networks in our results. These thresholds are in agreement with current recommendations for fMRI voxel- and cluster-based thresholding (Eklund et al. 2016).

## RESULTS

### Overview

Functional gradients 1 and 2 captured the highest portion of variability in DN resting-state functional connectivity to cerebral cortex (**Figure 1A**). Silhouette coefficient analysis identified three as the optimal number of k-means clusters when grouping DN voxels according to their functional gradient 1 and 2 values (**Figure 1C**). These three territories in DN corresponded uniquely to (i) default mode network; located rostral-ventrally in right and left DN, and also caudally in right DN; (ii) salience-motor network; located centrally in the rostral-caudal axis in right and left DN; and (iii) visual network; located caudally in left DN, and central-caudally in right DN (cluster locations within DN are shown in **Figure 2A**; functional connectivity patterns in cerebral cortex from each DN cluster are shown in **Figure 2B**). These functional connectivity patterns were successfully replicated in an independent dataset (**Figure 2B**).

**Figure 1.**
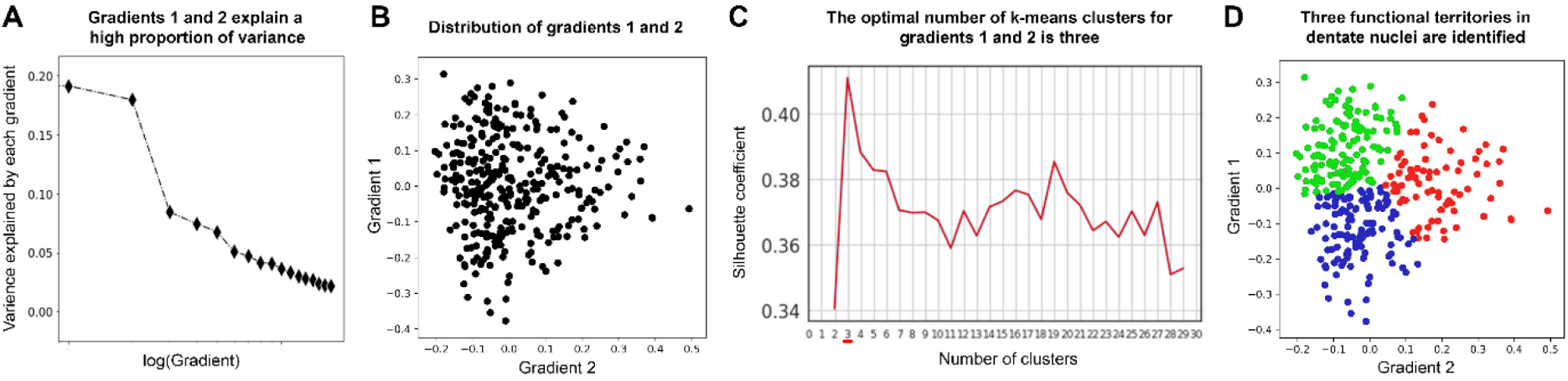
Data-driven identification of the number and location of functional territories in DN. **(A)** Functional gradients 1 and 2 capture the majority of variability in DN resting-state functional connectivity to cerebral cortex. Note a clear knee in the curve of variability explained by each functional gradient, indicating that the first two components are the most relevant aspects of DN functional neuroanatomy. **(B)** Distribution of DN functional gradients 1 and 2. Each dot in this graph corresponds to one voxel in left or right DN. **(C)** Silhouette coefficient analysis identified three as the optimal number of k-means clusters when grouping DN voxels according to their functional gradient 1 and 2 values. Note that the maximum score is achieved with three clusters. **(D)** K-means clustering grouped DN voxels in three categories according to their functional gradient 1 and 2 values. Each functional territory is shown here in a different color (red, blue, green), and these three colors correspond to the three functional territories shown in Figure 2A and 2B. See our online repository to access all custom code used for these analyses, https://github.com/xaviergp/dentate.

### Identification of the most relevant functional gradients of dentate nuclei

The first step to define the number and nature of functional territories in DN was to determine the amount of variance explained by each functional gradient. We observed a clear knee in the curve of data variability explained by each functional gradient, indicating that functional gradients 1 and 2 were the most relevant components of DN connectivity to cerebral cortex (**Figure 1A**). For this reason, only functional gradients 1 and 2 were included in further analyses. Clustering of voxels in DN based on functional gradients 1 and 2 values, and calculation of functional connectivity from each resulting cluster in our discovery and replication samples, provided a successful data-driven identification and characterization of DN functional territories, as follows.

### Identification of the optimal number of functional territories in dentate nuclei

As functional gradients 1 and 2 captured the highest portion of DN resting-state variability in our data, we considered that functional gradients 1 and 2 were an optimal dimensional space for the identification of functionally distinct territories in DN. A remaining question was the number of functional territories that our clustering analysis ought to identify. Unlike previous investigations that assumed a bi-modal motor versus non-motor division, we did not impose a-priori assumptions to determine the number of functional divisions with the DN. Instead, we conducted silhouette coefficient analysis and in this way determined that three was the optimal number of k-means clusters when grouping DN voxels according to their functional gradient 1 and 2 values (**Figure 1D**). We therefore concluded that three is the optimal number of functional subdivisions in human DN as indexed by high spatio-temporal resolution resting-state MRI in our sample.

### Clustering of three functional territories in the dentate nuclei

K-means calculation of three clusters was performed based on functional gradient 1 and 2 values (**Figure 1D**). This analysis required that all DN voxels contained within each functional territory were contiguous in functional gradient 1 and 2 coordinates (as shown in **Figure 1D**), but not necessarily contiguous in spatial coordinates. When grouped according to spatial localization in DN, multiple subclusters were identified for each of the three functional territories. Size and localization of each subcluster is reported in **Supplementary Table 1**. A minimum size of 10 contiguous voxels in spatial coordinates was imposed in order to eliminate noise in the spatial distribution of each functional territory. This resulted in the retention of three clusters for functional territory 1 (one in left DN located rostral-ventrally, one in right DN located rostral-ventrally, and one in right DN located caudally), two clusters for functional territory 2 (one in left DN and one in right DN, both located centrally along the rostral-caudal axis), and two clusters for functional territory 3 (one in left DN located caudally, and one in right DN extending from central to caudal territories along the rostral-caudal axis) (**Figure 2A**). Results of both thresholded (minimum size 10 voxels) and unthresholded functional territories are included in our online repository, but only thresholded maps are shown here.

**Supplementary Table 1.** For each functional territory, only subclusters of 10 or more voxels were retained for further analyses (shown in bold). This thresholding step decreased noise in the spatial distribution of the three functional territories in DN.

| FUNCTIONAL TERRITORY 1 |  |  | FUNCTIONAL TERRITORY 2 |  |  | FUNCTIONAL TERRITORY 3 |  |  |
| --- | --- | --- | --- | --- | --- | --- | --- | --- |
| Subcluster number | Number of voxels | Size (mm<sup>3</sup>) | Subcluster number | Number of voxels | Size (mm<sup>3</sup>) | Subcluster number | Number of voxels | Size (mm<sup>3</sup>) |
| **1** | **82** | **656** | **1** | **77** | **616** | **1** | **28** | **224** |
| **2** | **35** | **280** | **2** | **40** | **320** | **2** | **25** | **200** |
| 3 | 7 | 56 | 3 | 3 | 24 | **3** | **11** | **88** |
| 4 | 5 | 40 | 4 | 3 | 24 | 4 | 4 | 32 |
| 5 | 1 | 8 | 5 | 1 | 8 | 5 | 3 | 24 |
| 6 | 1 | 8 | 6 | 1 | 8 | 6 | 3 | 24 |
| 7 | 1 | 8 | 7 | 1 | 8 | 7 | 1 | 8 |
|  |  |  | 8 | 1 | 8 | 8 | 1 | 8 |
|  |  |  | 9 | 1 | 8 | 9 | 1 | 8 |
|  |  |  |  |  |  | 10 | 1 | 8 |
|  |  |  |  |  |  | 11 | 1 | 8 |

### Characterization of functional territories in dentate nuclei

Having identified the optimal number and spatial location of functional territories in DN, we aimed to characterize the functional contribution of each territory. This characterization was based on seed-to-voxel resting-state function connectivity analyses from each DN functional territory to cerebral cortex. This approach revealed three distinct functional connectivity patterns (**Figure 2B**), as follows: The first territory was functionally connected to the default-mode network, including connectivity to medial prefrontal cortex (mPFC), posterior cingulate cortex (PCC), angular gyrus (AG), and middle temporal lobe (MTL). The second territory corresponded to a salience-motor network, including connectivity to primary motor cortex (M1) and supplementary motor area (SMA), as well as insula (Ins), dorsal anterior cingulate cortex (dACC), anterior supramarginal gyrus (aSMG), and middle frontal gyrus (MFG). The third territory corresponded to a visual network, including connectivity to primary visual (V1) and visual association areas (VAA). These networks were observable when visualizing functional connectivity maps based on median 95^th^ percentile-thresholded r values in the discovery sample – namely, the same thresholds and sample that were used when calculating functional gradients (**Figure 2B**, first row). These networks were also observable when performing statistical significance thresholding (voxel and cluster-based p value thresholds) and calculating unique effects of connectivity from each DN functional territory (i.e., Functional Territory 1 > Functional Territory 2 + 3; Functional Territory 2 > Functional Territory 1 + 3; and Functional Territory 3 > Functional Territory 1 + 2) in the discovery sample (**Figure 2B**, second row). This analysis of unique effects is relevant as it provides statistical proof for the conclusion that connectivity patterns from each DN functional territory are different from the connectivity patterns of the other two DN functional territories in our sample. This conclusion would not be fully supported if connectivity effects from each functional territory were only analyzed independently. Importantly, similar patterns of connectivity were replicated in an independent sample (shown in two different statistical thresholds in **Figure 2B**, third and fourth rows). This replication analysis is relevant as it supports the extrapolation of these regions to future independent human studies of DN, providing reassurance that DN functional territories described here are not overfitted to one particular sample.

## DISCUSSION

Here we show for the first time that human DN is optimally divided into three functional territories as indexed by high spatio-temporal resolution resting-state MRI of 60 healthy participants, and that these three territories contribute uniquely to default-mode, salience-motor, and visual brain networks. Contrasting with previous views that define motor versus nonmotor territories as the principal poles of DN organization, the results presented here indicate that DN subdivisions span a broad range of human macro-scale neural specialization categories, namely default-mode, attentional, and multiple dimensions of unimodal processing including motor and visual. These conclusions are supported by the analytical strategies of 64-channel MRI imaging, data-driven methodology, and replication in an independent sample.

### Human dentate nuclei are divided into default-mode, salience-motor, and visual processing territories, echoing a broad spectrum of human macro-scale neural specialization

Each functional territory identified within the DN exhibited a unique pattern of functional connectivity with the cerebral cortex that corresponded to well-established brain networks (**Figure 2**). Here we argue that a broad spectrum of human macro-scale neural specialization is represented in DN, and that central components of this spectrum of functional diversity are segregated in distinct functional territories within DN.

Functional territory 1 corresponded to default-mode network (DMN), which we conceptualize here as the apex of the central axis of human macro-scale brain organization, as follows. Functions in the human brain can be classified according to multiple criteria. One fundamental classification method distinguishes sequential hierarchical levels of information processing – from primary processing cortices, to intramodality association areas, and progressively transmodal processing in heteromodal, paralimbic, and limbic cortices. This unimodal-to-transmodal progression is subserved by cerebral short association fibers that link primary processing cortices to progressively higher transmodal brain territories (Mesulam 1998, Pandya & Yeterian 1985). This fundamental axis of human brain organization can be captured in-vivo in cerebral cortex (Margulies et al. 2016), and it has been recently shown that cerebral-cerebellar anatomical connectivity results in a remarkably similar organization in cerebellar cortex (Guell et al. 2018b). Along this central unimodal-to-transmodal axis of brain specialization, DMN is situated at the highest transmodal extreme (Guell et al. 2018b, Margulies et al. 2016), with functional contributions revolving around unfocused cognitive processing such as mind wandering and related mental states of autobiographical memory retrieval and introspection (Buckner et al. 2008). All central nodes of DMN were included in cerebral cortical connectivity from functional territory 1, namely, PCC, mPFC, AG, and MTL (Buckner et al. 2008).

Functional territory 2 corresponded to salience-motor network, presented here as the cognitive opposite pole of default-mode network. Specifically, attentional functions in the brain are fundamentally distinct and arguably opposite to default-mode cognitive processes such as mind wandering. This view is supported across multiple human and animal studies of brain physiology. Attentional and DMN territories are anti-correlated at rest (Fox et al. 2005), dissociated in vigilant as opposed to inattentive brain state activity (Barch et al. 2013, Hayden et al. 2009), and causally interfere with the activation of each other (Chen et al. 2013). Within the realm of attention, cognitive neuroscience clearly delineates a fundamental distinction between goal-directed and stimulus-driven processes. Attention directed by top-down cognition is in clear contrast with attention dominated by external events (Corbetta & Shulman 2002). This distinction is also echoed in the intrinsic organization of brain functional architecture (Fox et al. 2006, Seeley et al. 2007) – such investigations have resulted in the definition of multiple attentional brain networks such as dorsal attention network and salience network. While all attentional systems are dissociated from DMN, salience network is uniquely segregated from DMN function because it is responsible for the sharp transition from inattentive/default-mode states to vigilant states (Goulden et al. 2014, Sridharan et al. 2008, Zhou et al. 2018). Thus in contrast with functional territory 1, functional territory 2 revealed functional connectivity to all central nodes of salience network, namely insula and dACC (Seeley et al. 2007), as well as secondary salience-linked territories including MFG and aSMG (see their connectivity to insula and dACC in Yeo et al. 2011 17-network parcellation).

Functional territory 2 also revealed connectivity to motor territories M1 and SMA, which we view here as a characterization of the nature of attentional processing in DN. It is well established that stimulus-driven attentional control is linked to sensorimotor systems, with anatomical studies revealing sensorimotor afferent and efferent projections in insula (Mesulam & Mufson 1982, Mufson & Mesulam 1982), and neuroimaging and invasive stimulation studies in humans confirming a physiological coupling between these two systems (Deen et al. 2011, Seeley et al. 2007, Stephani et al. 2011, Uddin 2015). A link between salience and motor processing in DN resonates not only with the well-established role of the cerebellum in motor control, but also with the anatomical and functional connections linking salience processing to sensorimotor processing in the brain.

Functional territory 3 corresponded to the visual network, presented here as the unimodal opposite pole of sensorimotor processing. Clustering of brain territories based on intrinsic functional connectivity patterns identifies somatosensory and visual as the two central unimodal components of brain organization (Yeo et al. 2011), and the transition from sensorimotor to visual functional connectivity patterns is the second largest source of variability in functional gradients of the cerebral cortex (Margulies et al. 2016). Functional territory 3 revealed connectivity to V1 and visual association areas, defining the third and last central division of human DN organization. Whereas connectivity to V1 was not observed in the replication sample analysis, visual association areas were present in both discovery and replication samples, supporting the validity of a visual functional territory in human DN. While there does not seem to be a predominant representation of primary visual processing in cerebellar cortex (Buckner et al. 2011), there are large territories of cerebellum devoted to attention processes relevant for visual association processing (Buckner et al. 2011, Guell et al. 2018b), and cerebro-cerebellar anatomical circuits include cerebral cortical visual association areas (Schmahmann & Pandya 1992, 1993). Moreover, recent evidence indicates that visual cognition is not only largely represented in cerebellum, but also organized in retinotopic maps that overlap with top-down attention networks in cerebellar cortex, specifically dorsal attention network (van Es et al. 2018). Visual association systems in DN may thus correspond predominantly to top-down attentional control, in contrast with salience-motor contributions of functional territory 2 that correspond predominantly to bottom-up attentional control.

### Default-mode, salience-motor, and visual processing domains are located in distinct regions within the dentate nuclei

Our results identified that central components of the spectrum of human neural functional specialization – default-mode, salience-motor, and visual – are segregated in distinct spatial territories within DN (**Figure 2A**). Default-mode processing was localized in rostral-ventral aspects of left and right DN, as well as in caudal aspects of right DN. Salience-motor processing was localized centrally along the caudal-rostral axis, and visual processing was localized caudally. This distribution of salience-motor and visual territories was more apparent in left compared to right DN. Specifically, there was more interdigitation between salience-motor and visual processing territories in right DN than in left DN, and the caudal portions of right DN were occupied both by default-mode and visual processing territories.

The broad functional domain categories identified here (default-mode, salience-motor, and visual) are different from previous descriptions of DN organization centered on a bi-modal motor versus non-motor division. For this reason, the anatomical location of our findings are not directly comparable to previous DN parcellations (Bernard et al. 2014, Dum & Strick 2003, Steele et al. 2017). However, our results align and in some cases contrast with some aspects of previous descriptions of DN functional neuroanatomy, as follows.

Our data-driven approach did not identify any territory of pure primary motor specialization in DN. This observation is in contrast with the well-established anatomical knowledge that non-human primate dorsal DN is predominantly linked to primary motor cortex (as reviewed in Dum & Strick 2003). It has been shown that ventral DN territories dedicated to nonmotor control are enlarged in human compared to lower species (Leiner et al. 1991, Matano 2001). Our findings support this possibility. We speculate that human DN territories specialized in pure motor control with connectivity only to sensorimotor territories may be too small, or may represent a too small portion of resting-state data variability, to be identified by our data-driven functional parcellation method as a separate component of human DN functional specialization. It is worth mentioning that the only previous investigation of resting-state human DN functional topography, based on manual resting-state seed location with the specific objective to identify pure motor versus pure nonmotor territories, did not successfully identify any DN territory with functional connectivity only to motor systems (Bernard et al. 2014, note that motor seed includes also connectivity to inferior parietal lobule and prefrontal cortex).

Current consensus in DN functional neuroanatomy is that ventral and caudal territories are specialized in nonmotor control, while dorsal territories are specialized in motor control. This bi-modal division was established in a compendium of anatomical investigations in monkeys by Dum and Strick (Dum & Strick 2003), and echoed in human investigations of resting-state connectivity (Bernard et al. 2014), task fMRI (Küper et al. 2011, 2012, 2014; Thürling et al. 2011), and tractography (Steele et al. 2017). All three principal subdivisions of DN organization identified in our study contained components of non-motor control. However, within non-motor domains, the prediction is that default-mode processing should be located preferentially in ventral territories of DN, as DMN is arguably the apex of a motor-to-nonmotor hierarchy of neural function (Guell et al. 2018b, Margulies et al. 2016, Mesulam 1998). Our findings are consistent with this prediction, as default-mode processing was the only functional territory with a spatial distribution that did not encroach upon dorsal aspects of DN. Specifically, both right and left DN revealed DMN processing that was segregated in rostral-ventral territories. Default-mode processing was also represented in right DN caudal pole. Perhaps consistent with this observation, anatomical connectivity reports map non-motor processing in the caudal pole of DN in addition to its ventral surface (Dum & Strick 2003), and task fMRI investigations also report non-motor processing in caudal aspects of DN (Küper et al. 2011, 2014; Thürling et al. 2011).

Representation of DMN in two territories in right DN (rostral-ventral and caudal), contrasting with one territory in left DN (rostral-ventral), is in agreement with whole-brain patterns of DMN lateralization. DMN is more predominantly present in left cerebral cortex (Agcaoglu et al. 2014) and right cerebellar cortex (Buckner et al. 2011). Right DN receives projections from right cerebellar cortex, and right cerebellar cortex is predominantly connected to left cerebral cortex. Stronger representation of DMN in the right DN is consistent with this knowledge. Left cerebral cortical and right cerebellar cortical language processing lateralization might also relate to our observed pattern of stronger lateralization of DMN to right DN. Of note, DMN territories overlap with language processing areas in cerebral and cerebellar cortex as indexed by some task fMRI paradigms (see supplementary material in Guell et al. 2018b). Cerebral cortical maps of connectivity identified here as DMN (**Figure 2**, functional territory 1) also included language-related regions such as angular gyrus and inferior frontal gyrus (Price 2012), and previous task fMRI experiments have demonstrated stronger language-related processing in right DN compared to left DN (Küper et al. 2011, Thürling et al. 2011). Further, the observation of multiple representations of DMN in right DN raises the larger possibility of network architecture within DN. It has long been established that numerous functional domains are represented in multiple segregated nodes in the cerebral cortex (Cavada & Goldman-Rakic 1989a,b; Selemon & Goldman-Rakic 1988, Yeo et al. 2011). Similarly, there are two representations of motor processing and three representations of multiple non-motor domains in cerebellar cortex (Guell et al. 2018a). The identification of two distinct nodes of DMN processing in right DN hints at the possibility that a multi-nodal representation of neural functions might also exist within DN.

**Figure 2.**
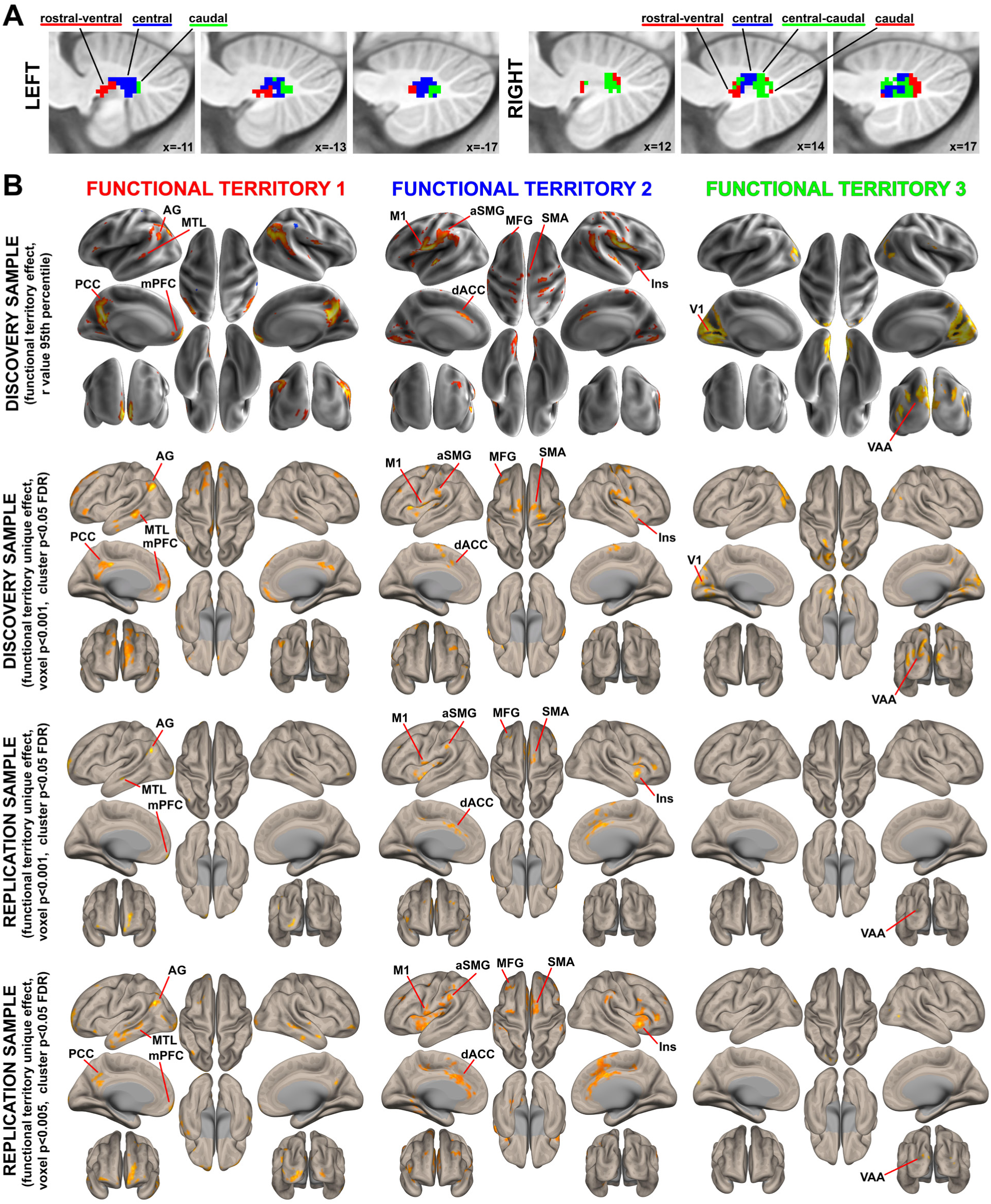
Spatial location and functional characterization of functional territories in DN. Green, red, and blue colors correspond to the same territories in Figures 1D, 2A, and 2B. **(A)** Spatial location of the three functional territories identified as shown in, Figure 1D. **(B)** Each functional territory identified in DN exhibited a unique functional connectivity pattern to cerebral cortex that corresponded to well-established brain networks. Specifically, functional territories 1, 2, and 3 corresponded to default-mode, salience-motor, and visual networks, respectively. First row: 95th percentile of median r values in the discovery cohort. Second row: statistical significance thresholding of the unique effect of each functional territory (e.g. functional territory 1 > functional territory 2 and 3), voxel p<0.001 with cluster size FDR correction of p<0.05. Note that calculations of unique effects are relevant as they provide statistical proof for the conclusion that functional connectivity from each functional territory in DN is different from the rest of territories. Third row: statistical significance thresholding equal to second row, but in an independent replication sample. Note that a replication analysis is important as it provides reassurance that functional territories reported here are not overfitted to the discovery sample. Fourth row: same analysis as third row, but with lower statistical thresholding to allow better visualization of brain networks (voxel p<0.005 with cluster size FDR correction of p<0.05). AG, angular gyrus; MTL, medial temporal lobe; PCC, posterior cingulate cortex; mPFC, medial prefrontal cortex; aSMG, anterior supramarginal gyrus; M1, primary motor cortex; MFG, middle frontal gyrus; SMA, supplementary motor area; Ins, insula; dACC, dorsal anterior cingulate cortex; V1, primary visual cortex; VAA, visual association area.

Visual processing was identified as the third central component of DN functional neuroanatomy. It included connectivity to primary and association visual areas, and was segregated in caudal and central-caudal components of left and right DN, respectively. Some previous reports are in agreement with a caudal distribution of visual processing in DN. Specifically, intracranial recording in monkeys revealed caudal DN neurons that responded specifically to “movement of the experimenter’s arm or pieces of paper moved in particular parts of the visual field” that could not be attributed to eye movements (van Kan et al. 1993, p. 62), with previous investigations also reporting selective light stimulus-related neurons in caudal DN (Chapman et al. 1986). More recently, task fMRI reports have identified visualspatial task activation in caudal DN bilaterally (Küper et al. 2011). It has also been demonstrated that non-human primate cortical visual inputs target specific territories of cerebellar cortex that then project to ventromedial aspects of DN (Xiong & Nagao 2002). Perhaps in agreement with the report of Xiong and colleagues, visual territories identified here in left and right DN encroached upon ventral DN surface.

### Relevance for cerebellar neuroscience, neurology, and psychiatry

Our results provide a systems-neuroscience substrate for cerebellar output to influence multiple broad categories of neural organization – namely default-mode, attentional-motor, and multiple unimodal streams of information processing including motor and visual. Brain networks such as salience and default-mode, here mapped to distinct DN territories for the first time, are central players in the evolving understanding of systems neuroscience in neurology and psychiatry. Identifying and mapping these macro-scale networks in different brain territories is relevant for the characterization of functional and structural abnormalities in psychopathology (Darby et al. 2018, Palaniyappan & Liddle 2012, Whitfield-Gabrieli & Ford 2012), and necessary for the development of targeted brain stimulation interventions (Esterman et al. 2017). In the specific case of cerebellar cortex, damage to the cerebellar posterior lobe, which provides inputs to the DN, causes the cerebellar cognitive affective syndrome, characterized by deficits in executive, linguistic, visuospatial and affective processing (Guell et al. 2015; Hoche et al. 2016, 2018; Schmahmann & Sherman 1998, Schmahmann et al. 2009, Stoodley et al. 2016). In addition, multiple studies have reported cerebellar cortical abnormalities in numerous disorders such as Alzheimer’s disease (Guo et al. 2016, Jacobs et al. 2018), autism spectrum disorder (Arnold Anteraper et al. 2018, D’Mello & Stoodley 2015), and schizophrenia (Moberget et al. 2018), with preliminary evidence suggesting that cerebellar cortical stimulation might improve symptoms in these diseases (Brady et al. 2019, Demirtas-Tatlidede et al. 2010, Di Lorenzo et al. 2013, Garg et al. 2016, Stoodley et al. 2017, Tikka et al. 2015). The importance of our findings is thus underscored by the relevance of cerebellar cortex in neurology and psychiatry (Schmahmann et al. 2019), as the majority of fibers exiting the cerebellar cortex synapse in DN before reaching extracerebellar structures such as cerebral cortex.

A better understanding of DN functional organization is relevant for the characterization of cerebellar and cerebellar-linked neuropathology. DN parcellation results (**Figure 2A**) made publicly available in our online repository provide new scientific knowledge concerning the functional organization of human DN, and the first data-driven functional parcellation of human DN. The replication of these findings suggests that these parcellations are generalizable to independent studies, with the cautionary note that future research is required to test the prediction that the same functional connectivity patterns will remain observable in adult populations, as discussed in the following section. Classical reports demonstrated topography of DN histological changes following lesions to distinct territories of the cerebral hemispheres, dating as back as 1927 (as reported in Smyth 1941). More recent reports indicate that specific neurological symptoms in cerebellar disease may correlate with atrophy in specific locations within DN (Ilg et al. 2013). The present observations provide an opportunity for an improved topographical interpretation of the functional significance of these findings. Similarly, previous investigations in psychiatry have observed abnormalities in functional connectivity when using all combined territories of DN as a seed (Olivito et al. 2017). Our results provide a more precise method of DN seed selection that might improve the detection of DN functional connectivity abnormalities in patient populations.

### Limitations and future studies

Functional characterization of territories in DN was based on resting-state connectivity with cerebral cortex. First, while the connectivity patterns observed here corresponded to well-established brain networks, future investigations of task-based fMRI may provide an improved characterization of the functional contributions of each DN compartment. Second, our investigation was focused on cerebral cortical patterns of connectivity, and did not analyze connectivity from DN to cerebellar cortex or other subcortical structures. One methodological limitation of analyzing connectivity to cerebellar cortex is that functional connectivity clusters corresponding to within-DN connectivity may bleed into cerebellar cortex given the close proximity of the two structures. Within other territories of subcortex, functional connectivity values are generally weaker as fMRI signal in the subcortex is generally worse than in the cerebral cortex. However, there are numerous anatomical projections to the DN from cerebellar cortex, and DN ascending projections are directed principally to the cerebral cortex after an obligatory synapse in the thalamus. Mapping connectivity from DN to cerebellar cortex and thalamus is thus an important area of functional neuroanatomy that could be further explored in future investigations. Third, resting-state functional connectivity is inherently limited due to its correlational nature – a causal demonstration of human DN functional territories could be achieved with modulation/stimulation experiments. Recent developments utilizing these methods suggest that it might be possible in the near future to target specific sub-territories in human DN non-invasively (Folloni et al. 2019, Grossman et al. 2017, Lee et al. 2015, 2016; Verhagen et al. 2019). Of note, mean age was 13.64 years in the discovery sample and 15.13 years in the replication sample. Future work is required to test the prediction that our characterization of DN functional architecture will remain consistent in adult populations. Importantly, studies in this topic indicate that basic network organization – what the principal brain functional networks are (DMN, attentional, primary), and their position in the brain – does not change from adolescence to adulthood (see first figure in Marek et al. 2015, comparing network organization in childhood, early adolescence, late adolescence, and adulthood).

Our study reveals for the first time that DN subdivisions span a broad spectrum of human macro-scale neural specialization categories, namely default-mode, attentional, and multiple dimensions of unimodal processing including motor and visual. This observation is consistent with the anatomical knowledge that large territories of cerebellar cortex project to DN. As cerebellar cortex contains representations of default-mode, attentional, and multiple unimodal domains, it is logical to detect a similar spectrum of functional diversity in DN. This reasoning leads us to hypothesize that a similar organization should be observable in analogous brain compartments – such as pontine nuclei, that are an obligatory relay of topographically arranged cerebral cortical projections reaching the cerebellar cortex (Schmahmann 1996; Schmahmann & Pandya 1995, 1997, Schmahmann et al. 2004a,b), or inferior olivary nuclei, that send and receive projections to and from cerebellar nuclei in a topographically precise fashion (Holmes & Stewart 1908, Voogd et al. 2013). In this way, DN findings presented here unmask new predictions for basic human functional neuroanatomy of other largely unexplored brain territories.

## ACKNOWLEDGEMENTS

This work was supported in part by La Caixa Banking Foundation (100010434, LCF/BQ/AN15/10380048 to X.G.); the MGH Tosteson & Fund for Medical Discovery Award (to X.G.); US National Institutes of Health (NIH) grants 1U01NS104326-01 (to J.D.S.), 1R01NS080816-01A1 (to J.D.S.), U01MH108168 (to J.D.E.G. and S.W.G.), F32MH114525 (to N.A.H.), F32MH117933 (to A.M.D.), T32MH112510 (to R.R.R.), and F31HD086957 (to R.R.R.); the William and Flora Hewlett Foundation (4429) (to J.D.E.G.); the Simons Center for the Social Brain (postdoctoral fellowship to A.M.D.); the National Ataxia Foundation (to J.D.S.); the MINDlink foundation (to J.D.S.); and the Athinoula A. Martinos Imaging Center at the McGovern Institute for Brain Research at MIT. The authors thank Scott Marek for his help reviewing the literature regarding differences between adolescent and adult brain functional architecture; Viviana Siless, Anastasia Yendiki, Stefan Hofmann, Diego Pizzagalli, Randy Auerbach, and Aude Henin for their contributions to the U01MH108168 study; and Yoon Ji Lee for technical assistance with data compilation.

